# A pectin-based formulation protects milk fat globule membranes in stored human milk

**DOI:** 10.64898/2026.07.13.734508

**Authors:** Justin E. Silpe, Hyesoo Kim, Anjali ShahLyng, Yi-Ting Tsai, Kelsey E. Johnson, Bum Jin Kim, Carolyn M. Slupsky, Ameer Y. Taha, David C. Dallas, Itay Budin, Bonnie L. Bassler

## Abstract

During household storage, expressed human milk can develop odor and flavor changes that trigger infant refusal and lead caregivers to discard their saved milk supply. We show that typical refrigeration and freezing conditions disrupt the milk fat globule membrane (MFGM), exposing milk lipids to lipases that catalyze hydrolysis and oxidation. A pectin-based formulation (PBF) maintains MFGM integrity during storage and following lipase challenge, suppressing production of glycerol, free fatty acid, and oxylipin byproducts without broadly affecting milk macronutrients, the proteome, and culturable microbial burden. Across an independent cohort of lactating individuals, lipase activity varied but tracked with maternal milk lipase gene expression, implicating endogenous lipolysis in stored-milk deterioration. In a blinded olfactory panel, PBF-treated, lipase-challenged milk smelled more like fresh milk than untreated controls. Together, these findings show that stabilizing the MFGM can protect stored human milk from lipase-driven deterioration, preserve sensory quality, and support use for infant feeding.

## Introduction

Human milk provides the optimal source of nutrients for most infants, supporting healthy growth and immune development. Global health organizations recommend exclusive breastfeeding for the first 6 months of life, with continued breastfeeding for at least 12 months^1^. Despite these recommendations, breastfeeding rates in the United States fall far short of these goals. While 85.7% of infants are breastfed initially, only 27.9% are breastfed exclusively through 6 months, and only 40.8% receive any human milk by 12 months (CDC NIS-Child breastfeeding rates, https://www.cdc.gov/breastfeeding-data/survey/results.html; accessed 30 Apr 2026). These declines are largely due to challenges faced by lactating parents, including engorgement, latch difficulties, and return to work^2^. Pumping and storing human milk offer a practical strategy to overcome these challenges while allowing parents to maintain adequate supplies for continued human milk feeding. A recent national survey indicates that approximately 80% of respondents begin pumping milk within one week postpartum, and 70% of respondents maintain a regular pumping routine or schedule in the United States^2^. However, many infants reject stored human milk. Surveys report that ∼25% of infants refuse stored human milk, often in association with changes in smell or taste^3,4^. One common cause of rejection is rancidification, an unpleasant change in milk smell and taste resulting from lipid hydrolysis (i.e., lipolysis) or oxidation. Freezer storage is widely used to extend human milk shelf life, yet rancidification remains a concern, as it progresses with prolonged storage time under standard household freezing conditions^5^. As pumping and storing human milk have become increasingly common, discarding milk due to rancidification can be a significant and distressing experience for lactating parents^2,5^. Thus, strategies to halt or slow spontaneous lipid deterioration during human milk storage are urgently needed.

Lipase activity has been identified as a major driver of rancidification in stored human milk samples. Human milk contains two major lipases: bile salt-stimulated lipase (BSSL; encoded by *CEL*) and lipoprotein lipase (LPL; encoded by *LPL*), which facilitate breakdown of milk fat to support infant digestion^6^. Approximately 98% of total milk fat is composed of triacylglycerides (TAGs)^7^, a class of nonpolar lipids consisting of a glycerol backbone esterified to three fatty acids. TAGs serve as the primary substrate for lipolysis. Lipase-mediated TAG hydrolysis converts them into glycerol and free fatty acids (FFAs). Notably, human milk can undergo lipolysis when stored above −70 °C^8^, a temperature well below what typical home freezers can achieve, making lipase activity a persistent concern during household storage.

The remaining 2% fraction of milk fat consists of polar lipids, including phospholipids, sphingolipids, and cholesterol, that assemble into a unique trilayer structure known as the milk fat globule membrane (MFGM) surrounding the TAG core. The MFGM forms through a stepwise process during milk secretion from mammary epithelial cells^9^: An initial phospholipid monolayer enveloping the TAG core is derived from the endoplasmic reticulum during MFG synthesis, followed by acquisition of an outer bilayer from the apical plasma membrane during secretion into the alveolar lumen^10^ (Figure 1A). This trilayer membrane architecture enables the MFGM to function as a natural barrier that limits lipase access to the TAG core^11^. When the MFGM is disrupted, lipases can hydrolyze TAGs, releasing FFAs which generate the off-flavors that contribute to rancidification in stored human milk^12^.

**Figure 1.**
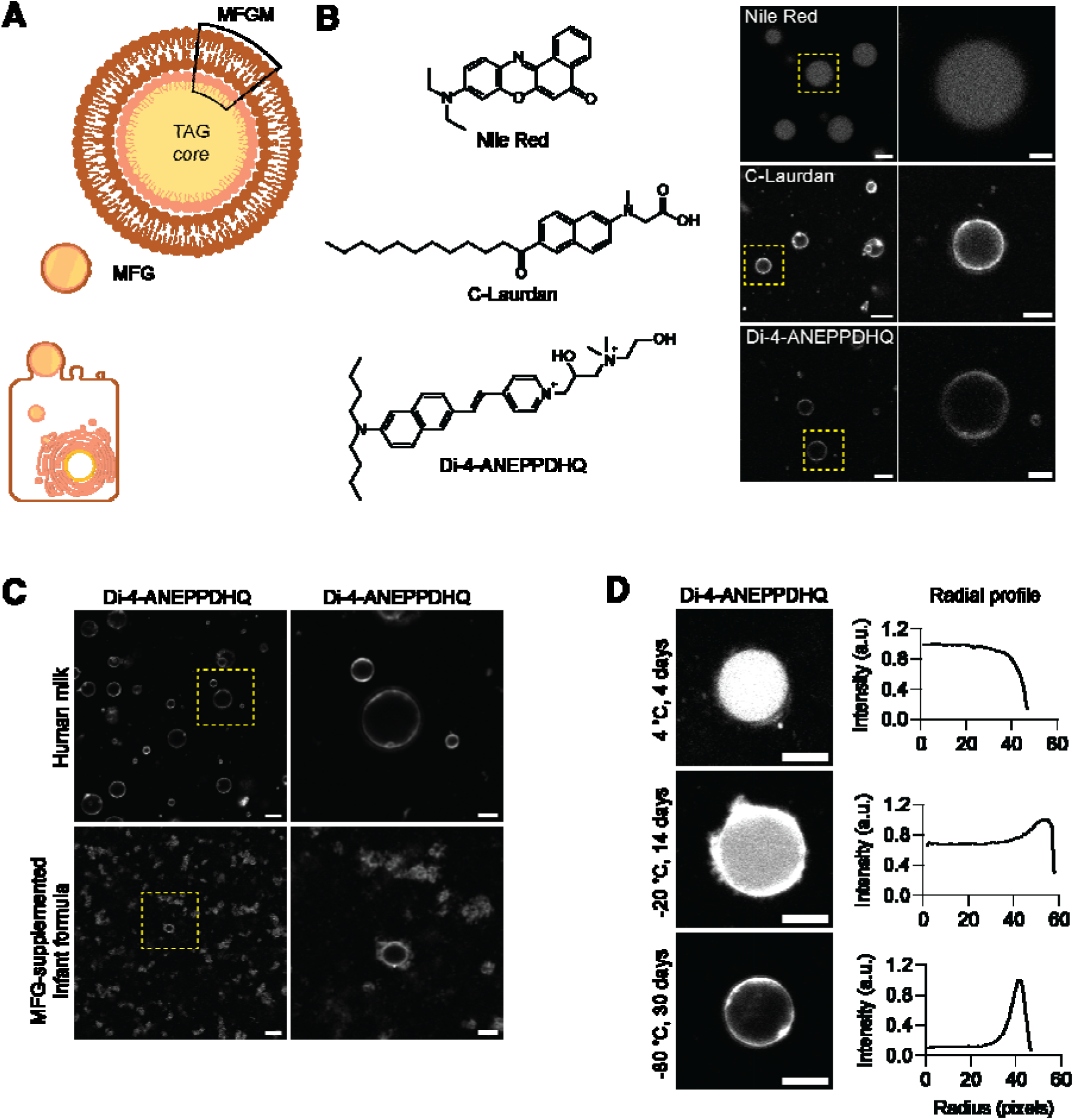
Di-4-ANEPPDHQ imaging of MFGM integrity. (A) Schematic of plasma membrane generating the MFG with its TAG core and trilayered MFGM. (B) Confocal micrographs of human milk MFGs stained with Nile Red, C-Laurdan, and Di-4-ANEPPDHQ. Yellow boxes indicate the enlarged regions shown in the right panels. Scale bars: 5 µm (left), 2 µm (right). (C) Di-4-ANEPPDHQ staining of human milk and MFGM-supplemented commercial infant formula. Yellow boxes indicate the enlarged regions shown in the right panels. Scale bars: 10 µm (left), 5 µm (right). (D) Di-4-ANEPPDHQ staining of human milk stored as designated. Scale bar: 5 µm. Radial fluorescence intensity profiles of individual MFGs shown to the right. Radial intensity was normalized to the maximum intensity of each MFG. Radius = 0 corresponds to the MFG core.

Beyond contributing to rancid flavor, lipolysis-generated FFAs also increase susceptibility to lipid oxidation^13^. Lipid oxidation occurs when unsaturated fatty acids react with oxygen, producing primary oxidation products called hydroperoxides, which further decompose into secondary products including aldehydes and ketones that are responsible for metallic off-flavors^14,15,16^. Lipid oxidation also generates reactive carbonyl species and radicals that promote glycation of human milk proteins and contribute to the formation of advanced glycation end products (AGEs) associated with inflammation and metabolic diseases^17,18^, further reducing the nutritional quality of stored human milk. Human milk contains endogenous antioxidants, including vitamins C and E, that help protect against lipid oxidation. However, vitamin C is rapidly depleted during storage^19^, and vitamin E levels can also decline under certain storage conditions^20^, progressively reducing antioxidant capacity of human milk over time.

These observations raise a practical question: can a food-based stabilizer preserve the lipid interface in expressed human milk during storage? Repurposing food-grade ingredients already approved for use in infant nutrition may offer a safe and practical intervention. We recently reported that a formulation combining pectin and ascorbic acid (vitamin C) preserves the sensory and lipid quality of stored human milk^21^. While the antioxidant roles of vitamins C and E in limiting lipid oxidation are well established^22^, the mechanism by which pectin reduced lipolysis was not explored. Understanding this mechanism is important for developing milk stabilizers and for defining the causes of stored-milk rancidification and infant refusal.

Here, we investigate this pectin-based formulation (PBF) as a human milk stabilizer. The PBF consists of low-methoxy pectin supplemented with vitamins C and E and is added to human milk at a final concentration of 0.2% (w/v) pectin. We apply the fluorescent dye Di-4-ANEPPDHQ to assess human milk MFGM integrity and quantify structural changes that occur under different storage conditions and PBF treatments. We test whether lipase activity impairs MFGM structure and whether PBF treatment mitigates lipase-induced MFGM disruption. We evaluate the relationship between *CEL* and *LPL* transcript abundance and net lipase activity by combining RNA-seq with lipase activity assays. Proteomic, metabolomic, and targeted oxylipin measurements, together with bulk milk composition and microbial burden measurements, are used to evaluate whether PBF treatment selectively protects the MFGM without broadly altering milk composition. Because premium infant formulas are often supplemented with bovine-derived MFGM ingredients to better approximate human milk composition, we also compare MFG structures between human milk and a commercial MFGM-supplemented formula to assess how such infant formula MFGM additives compare with the native MFGM architecture. Finally, we use a blinded olfactory assay to assess whether PBF treatment preserves sensory quality in a lipase-driven deterioration model.

## Results

### A cationic lipophilic dye enables visualization of MFGM integrity during human milk storage

Fluorescent membrane dyes offer non-destructive means to assess lipid membrane structure. For example, Nile Red has been widely used to stain neutral storage lipids commonly found in lipid droplets and MFGs^23^. However, tools that accurately report on MFGM integrity are not broadly known. To identify a suitable probe for visualizing the MFGM, we screened lipophilic dyes often used to stain cell membranes: Nile Red, C-Laurdan, and Di-4-ANEPPDHQ. Solvatochromic dyes such as C-Laurdan and Di-4-ANEPPDHQ commonly report on membrane organization through shifts in their fluorescence emission profiles that reflect lipid packing and membrane hydration^24^. Nile Red robustly stained the neutral TAG core but failed to resolve the surrounding membrane structure, whereas C-Laurdan preferentially labeled the MFGM but photobleached rapidly under our imaging conditions (Figure 1B). By contrast, Di-4-ANEPPDHQ produced a stable peripheral fluorescence signal that enabled clear visualization of the MFGM. The superior specificity of Di-4-ANEPPDHQ compared to Nile Red is likely due to the amphipathic and cationic structure of Di-4-ANEPPDHQ, which promotes partitioning into the outer leaflet of the MFGM and prevents rapid flip-flopping to the hydrophobic TAG core. To our knowledge, this study represents the first application of Di-4-ANEPPDHQ to probe MFGM integrity in human milk.

Using Di-4-ANEPPDHQ, we compared MFG structures in human milk and MFGM-supplemented infant formula (Figure 1C). Human milk possessed large, well-defined spherical MFGs surrounded by distinct membrane fluorescence. MFG structures in MFGM-supplemented formula appeared smaller and were frequently aggregated. Together, these images demonstrate that Di-4-ANEPPDHQ reports on intact MFGM organization in human milk, and this comparison shows that MFGM-supplemented infant formula does not reproduce the large, membrane-bound MFG architecture present in human milk.

Having established Di-4-ANEPPDHQ as a robust reporter of MFGM structure, we next examined whether it could detect changes in MFGM integrity arising from different storage conditions (Figure 1D). Human milk stored at −80 °C for 30 days exhibited Di-4-ANEPPDHQ fluorescence confined to the MFGM, consistent with an intact membrane structure. In contrast, human milk stored at 4 °C for 4 days showed loss of discernible MFGM staining, with no clear distinction between MFGM and TAG core fluorescence intensity. Human milk stored at −20 °C for 14 days displayed partially penetrated dye staining with increased fluorescence near the TAG core, indicative of compromised MFGM integrity relative to −80 °C storage. Radial fluorescence intensity profiles of individual MFGs confirmed these observations, with intact MFGs exhibiting peak fluorescence at the membrane and compromised MFGs showing increased signal toward the MFG center. Collectively, these results demonstrate that Di-4-ANEPPDHQ sensitively reports storage-dependent changes in MFGM integrity and provides an imaging-based readout of membrane stability during milk storage (Figure 1D).

### A pectin-based formulation preserves MFGM integrity during milk storage

Our prior high-throughput screen and storage study suggested that a PBF might slow lipid breakdown by preserving the MFGM. Here, we examined whether the PBF could maintain MFGM integrity under different storage conditions. Human milk samples frozen at −80 °C immediately after expression were thawed and then stored at 4 °C or −20 °C for defined intervals, with untreated and PBF-treated samples analyzed in parallel by Di-4-ANEPPDHQ imaging.

After 4 days at 4 °C, untreated samples showed marked loss of membrane-associated Di-4-ANEPPDHQ signal, and the distinction between the MFGM and TAG core was largely lost (Figure 2A). In contrast, PBF-treated samples retained peripheral Di-4-ANEPPDHQ signal, and radial-profile quantitation showed lower core-normalized dye penetration in PBF-treated aliquots from two of three donor samples, with no significant donor-level treated-versus-untreated differences (Figure S1). During frozen storage, untreated samples exhibited membrane-penetrant Di-4-ANEPPDHQ staining with increased dye access to the TAG core after 30 days at −20 °C, whereas PBF-treated samples maintained predominantly membrane-localized Di-4-ANEPPDHQ, closely mimicking samples stored at −80 °C (Figure 2B). Consistent with these images, radial-profile quantitation of PBF-treated aliquots from all three donor samples after 30 days at −20 °C showed lower core-normalized dye penetration than that in untreated samples (Figure S1).

**Figure 2.**
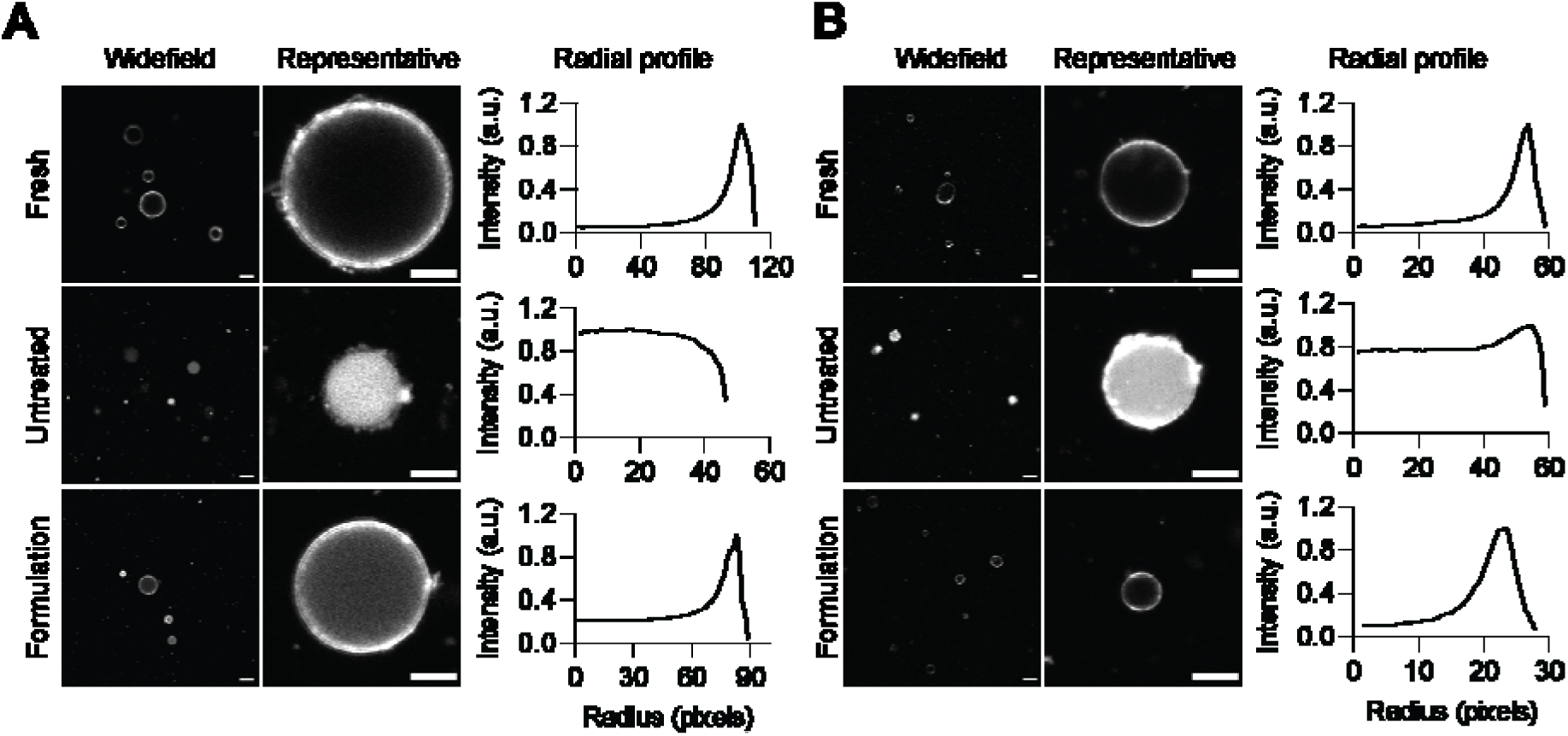
Di-4-ANEPPDHQ staining of untreated and PBF-treated milk during storage. (A) Widefield and magnified images after 4 days at 4 °C with or without PBF. Scale bars: 10 µm (widefield, left images), 5 µm (magnified, right images). (B) Images after 30 days at −20 °C with or without PBF treatment. Scale bars: 10 µm (widefield, left images), 5 µm (magnified, right images). In A and B, one representative MFG is shown for each widefield image, and its radial fluorescence intensity profile is provided to the right. Radial intensity was normalized to the maximum intensity of each MFG. Radius = 0 corresponds to the MFG core.

### The pectin-based formulation preserves MFGM integrity following exogenous lipase treatment

The above findings support a model in which MFGM destabilization increases lipase access to the TAG core. To test this supposition, we examined whether lipase addition destabilizes the MFGM and whether the PBF could preserve MFGM integrity following lipase treatment. Because human milk contains maternally-encoded lipases and could, in principle, also harbor bacterial lipases encoded by human milk-associated members of the microbiota, we examined both mammalian and bacterial lipases by time-lapse Di-4-ANEPPDHQ imaging.

Exogenous addition of mammalian lipase caused progressive disruption of untreated MFGMs, marked by loss of membrane-localized Di-4-ANEPPDHQ signal and increasing dye penetration into the TAG cores (Figure 3A). By 20 minutes, most MFGs within the field of view had lost discernible membrane structure, and by 30 minutes this loss was observed throughout the field. In PBF-treated samples, addition of mammalian lipase caused the Di-4-ANEPPDHQ to redistribute from the MFGM toward the TAG core, but compared to the untreated samples, membrane disruption was substantially attenuated. Indeed, despite some dye penetration, treated MFGs retained well-defined peripheral Di-4-ANEPPDHQ signal, consistent with preservation of MFGM structure during mammalian lipase exposure (Figure 3A).

**Figure 3.**
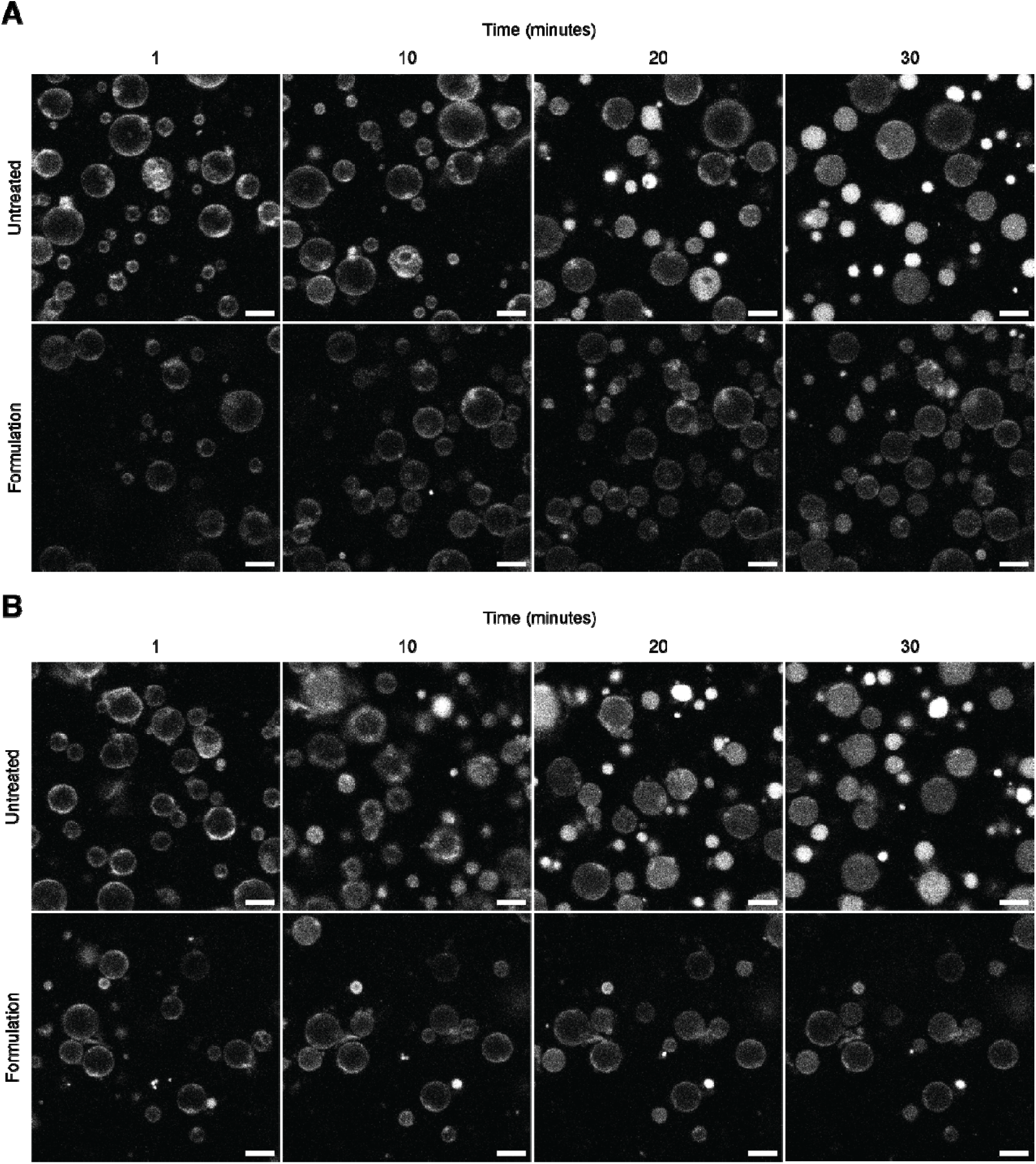
Time-lapse of Di-4-ANEPPDHQ staining of MFGs following administration of lipase. (A) 4 U/mL mammalian lipase treatment of untreated and PBF-treated milk. (B) 0.02 U/mL bacterial lipase treatment of untreated and PBF-treated milk. Times as designated. Scale bar: 5 µm.

Bacterial lipase acted more rapidly on the untreated MFG samples and at even lower dosages than the mammalian lipase, producing uniform dye distribution throughout the MFG and loss of discernible membrane structure by 10 minutes (Figure 3B). Strikingly, PBF-treated samples retained strong MFGM-associated Di-4-ANEPPDHQ signal through 30 minutes, indicating that the PBF delayed bacterial lipase-induced MFGM disruption.

### Natural variation in CEL and LPL transcript abundance tracks with net lipase activity in human milk

Because human milk lipase activity varies across donors and these differences could contribute to donor-specific extents of MFGM degradation during storage, we investigated whether variations in lipase activity were reflected at the level of expression of the genes encoding lipase enzymes. To position the storage phenotypes in the context of natural donor-to-donor lipase variation, we measured lipase activity in human milk samples collected at 1 and 6 months postpartum from an independent Mothers and Infants LinKed for Healthy Growth (MILK) Study cohort and compared the values we obtained with RNA-seq profiles generated from the same samples. Lipase activity varied substantially across samples. Regarding the two primary endogenous lipases present in human milk, transcript abundances for both *CEL* and *LPL* significantly correlated with measured lipase activity. Specifically, *CEL* and *LPL* transcript abundances showed positive associations (est = 0.33, P = 0.0012; Figure S2A and est = 0.30, P = 0.022; Figure S2B, respectively) with 4-nitrophenyl octanoate (4-NPO)-measured lipase activity. These data indicate that donor-to-donor differences in measured lipase activity are reflected at the transcript level for both major human milk lipases. We also visualized endogenous lipase activity directly in human milk: untreated human milk showed strong signal from a BODIPY-C12-based fluorogenic lipase substrate, whereas scalding at 82 °C for 10 minutes or treatment with orlistat, a lipase inhibitor, reduced the signal (Figure S3).

### Proteomics and bulk milk composition remain broadly stable during storage following pectin-based formulation treatment

While we observed that the PBF preserves MFGM integrity, a concern was whether the effect is specific to membrane integrity or reflects broader changes in milk composition. To assess specificity of the treatment, we profiled untreated and PBF-treated samples of a matched three-donor storage subset by liquid chromatography-tandem mass spectrometry proteomics at four storage timepoints (4 days at 4 °C, 7 days at −20 °C, 30 days at −20 °C, and 90 days at −20 °C).

Total protein profiles were highly similar between matched treatment groups (Pearson r = 0.98), with 185 to 190 proteins consistently identified across conditions and no broad PBF-associated shifts in relative abundance among major nutritional and immune proteins (Figure S4). Consistent with that unchanging protein profile, Fourier transform infrared (FTIR)-based bulk composition profiling, which provides a readout of macronutrient content, revealed no substantial PBF-associated shifts in milk fat, protein, or lactose across additional donor cohorts. Likewise, microbial plate counting showed no significant increase in culturable microbial burden following PBF addition (Figures S5 and S6).

### Targeted oxylipin and metabolite measurements reveal PBF treatment reduces lipid oxidation and lipolysis in stored human milk

Having found no evidence for general PBF-associated changes in the proteome or bulk milk composition of stored milk, we predicted that targeted biochemical measurements could reveal key PBF-driven differences in lipid deterioration during storage. We first focused on free oxylipins, early products formed by oxidation of free fatty acids. Importantly, bulk assays such as those for thiobarbituric acid reactive substances or measurements of hydroperoxide do not detect early oxidative changes within the lipid fraction. Thus, such bulk assays can only be reliably used to measure downstream oxidation products^16^. Because linoleic acid (18:2) is the most abundant polyunsaturated fatty acid (PUFA) in human milk, we focused on linoleic acid-derived oxylipins. The relevant derivatives are free hydroxyoctadecadienoic acids (HODEs), including 13-HODE, 9-HODE and free 12(13)-epoxy-9Z-octadecenoic acid (EpOME). Free 13-HODE and free 9-HODE were lower in PBF-treated milk than in matched untreated milk in all 12 non-baseline donor-condition comparisons. Free 13-HODE was 1.4-to 2.6-fold lower across stored timepoints, while free 9-HODE was 2.6-to 2.9-fold lower through 30 days, with a smaller, non-significant difference at 90 days. Free EpOME was lower in 11 of 12 comparisons and showed its strongest reduction after 4 days at 4 °C (8.6-fold; Figure 4).

**Figure 4.**
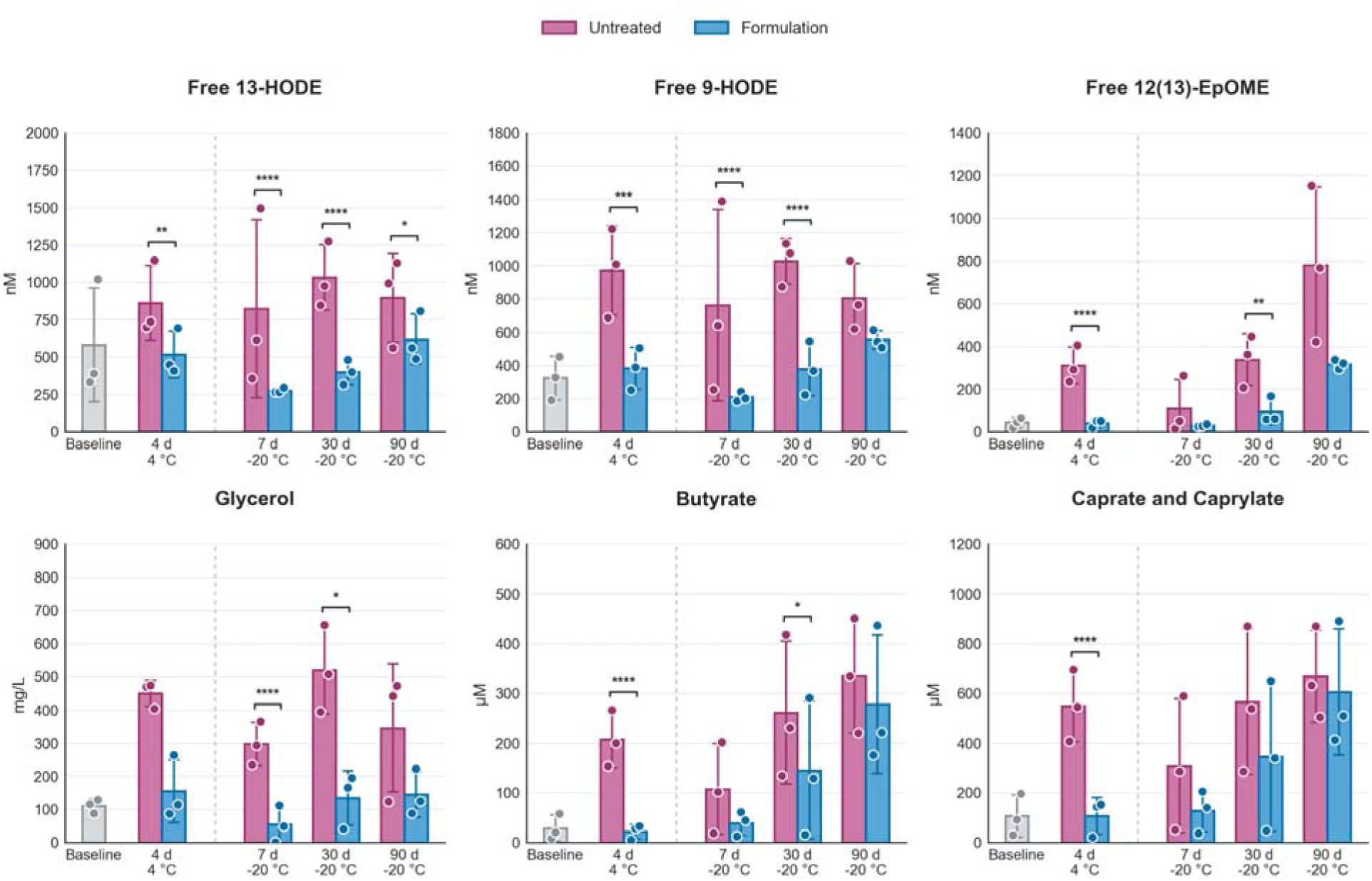
Quantitation of the levels of select compounds in untreated and PBF-treated human milk across storage conditions. Shown are concentrations of the designated compounds in untreated (magenta) and PBF-treated (blue) samples. Bars represent means across donors and error bars show SDs. Points show individual donor values. Baseline represents the untreated day-0 sample. The vertical dashed line separates the 4 °C and −20 °C storage conditions. Brackets indicate Holm-adjusted treated-versus-untreated contrasts from a linear mixed-effects model. * P < 0.05, ** P < 0.01, *** P < 0.001, **** P < 0.0001.

We next assessed whether treatment of stored milk with the PBF also reduced hydrolytic lipid breakdown products associated with rancid flavor. Glycerol, a direct product of TAG hydrolysis, was lower in PBF-treated human milk than in matched untreated milk in all 12 non-baseline donor-condition comparisons, with the strongest reduction after 7 days at −20 °C (16.8-fold lower, Figure 4). Short-chain and medium-chain FFA markers associated with rancid flavors in milk were also lower in PBF-treated samples. The largest reductions in these FFA markers occurred after 4 days at 4 °C, where butyrate and the combined caprate and caprylate signal were 12.1-fold and 7.0-fold lower, respectively, in PBF-treated human milk than in matched untreated milk. Indeed, across all non-baseline donor-condition comparisons, butyrate and the combined caprate and caprylate signal were lower in 11 of 12 and 10 of 12 matched comparisons, respectively (Figure 4).

In summary, these results indicate that PBF treatment does not measurably perturb the proteome nor bulk macronutrient composition of stored human milk; rather, it selectively suppresses oxidation-associated free oxylipins and hydrolysis-associated lipid breakdown products.

### Olfactory evaluation

Having measured lower accumulation of lipolysis and oxidation products in PBF-treated stored human milk than in untreated stored human milk, we next addressed whether these biochemical differences translated into a perceptible sensory outcome. To do this, we conducted a blinded, forced-choice olfactory evaluation using microbial lipase-treated goat milk as a surrogate mammalian system with and without PBF administration. Of 44 adult panelists, 36 (82%) identified the PBF-treated sample as more similar in odor to a fresh reference sample than the lipase-only-treated sample (binomial test, P < 0.001; Figure 5A). Panelists who selected the PBF-treated sample also rated it as more similar to fresh (median perceived similarity 75%) than those who selected the lipase-only sample (median 25%; Mann–Whitney U, P = 0.001; Figure 5B). These results indicate that PBF-mediated suppression of lipase-driven deterioration products is detectable by human olfaction.

**Figure 5.**
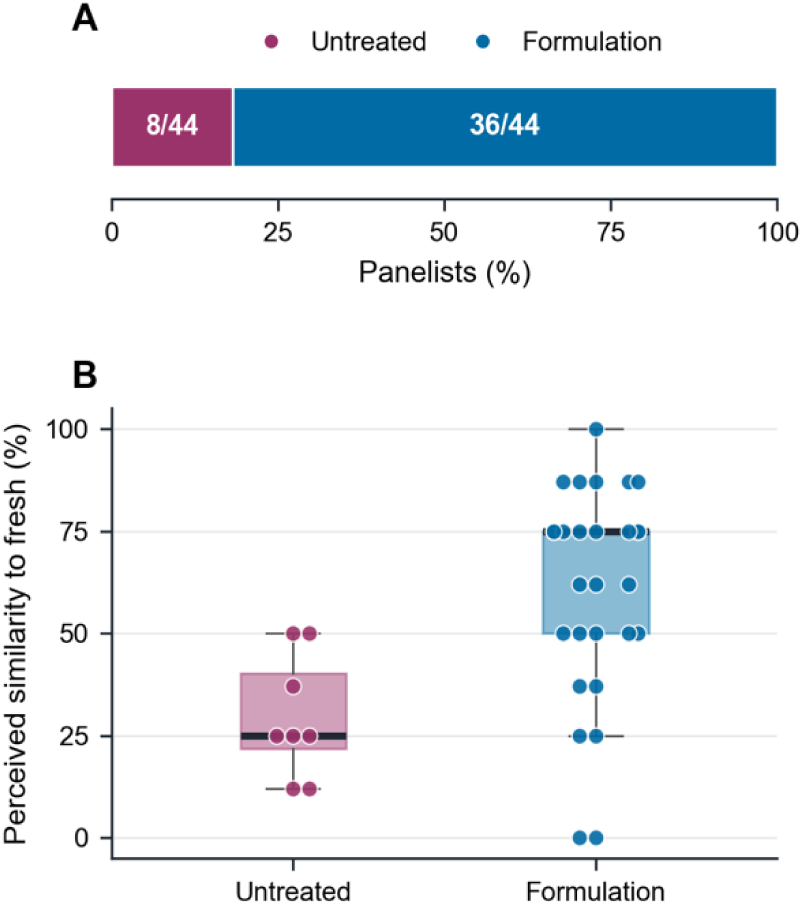
Blinded olfactory evaluation of PBF-treated versus lipase-only-treated goat milk. (A) Proportion of adult panelists (n = 44) selecting each sample as smelling more similar to a fresh reference in a forced-choice paradigm. (B) Perceived similarity to the fresh reference for each panelist’s chosen sample, rated on a 0-100 scale. In panel B, box plots show medians and interquartile ranges; points represent individual panelists.

## Discussion

Human milk MFGs consist of a neutral lipid core surrounded by a tri-layered MFGM enriched in glycans, bioactive proteins, and polar lipids that stabilize the emulsion and restrict enzymatic access to the lipid core prior to infant digestion. In our study, we identify disruption of the MFGM as a key feature of storage-associated deterioration of human milk, and we show that a PBF preserves MFGM integrity by limiting lipolysis and oxidation.

Using Di-4-ANEPPDHQ as a reporter of MFGM organization, we demonstrated that storage of human milk induces progressive disruption of the MFGM, reflected by redistribution of the dye from the membrane to the TAG core. This transition occurs within 4 days at 4 °C, consistent with current storage recommendations, and within fewer than 30 days at −20 °C. The latter likely reflects freeze-induced mechanical damage, such as ice crystal formation^25^, which compromises MFGM structure, and continued endogenous lipase activity, consistent with prior evidence that human milk can undergo lipolysis above −70 °C^8^. The difference between the consequences of −20 °C and −80 °C storage may reflect a combination of higher physical stress at household-freezer temperature and stronger suppression of residual lipase activity at ultralow temperature. Freeze-thaw cycling during routine household use could further exacerbate this difference. Thus, storage accelerates MFG deterioration, and importantly, well within current storage guidelines for consumption within 6-12 months in the freezer (CDC breast milk storage guidance, https://www.cdc.gov/breastfeeding/breast-milk-preparation-and-storage/handling-breastmilk.html; accessed 30 Jun 2026). Our imaging data show that adding the PBF preserves MFGM integrity across different storage conditions. At both 4 °C and −20 °C, the Di-4-ANEPPDHQ remained restricted to the peripheral membrane in PBF-treated samples, whereas untreated milk showed staining in the core, indicative of MFGM deterioration.

Loss of MFGM integrity is consistent with the known structural role of MFGM lipids. The MFGM outer layer is derived from the plasma membrane, which is particularly enriched in sphingolipids^26^. Sphingolipids contain saturated hydrocarbon chains that pack tightly with cholesterol to form ordered membrane domains that confer structural rigidity that protects the lipid core from lipase activity^27,28,29^. Consistent with this idea, sphingomyelin, the predominant sphingolipid in human and bovine milk, has been shown to slow lipolysis and fat absorption *in vivo*^30^, suggesting that sphingomyelin-rich MFGM domains contribute to restricting lipase access to the TAG core. The maintenance of MFGM-associated staining across different storage conditions in PBF-treated samples raises the possibility that pectin preferentially stabilizes sphingolipid-enriched outer leaflet of the MFGM.

MFGM integrity is closely linked to lipolysis, as lipases act on the TAG core once the membrane barrier is disrupted. Indeed, our study shows that pectin shields MFGs from lipases, including from pancreatic and bacterial lipases, by preserving the MFGM structure. When milk is expressed and stored, two endogenous lipases, BSSL and LPL, remain active above −70 °C^6,8^. Regarding these endogenous lipases, transcript abundances of both *CEL*, which encodes BSSL, and *LPL* correlate with measured lipase activity (Figure S2A and Figure S2B). Microscopy with a BODIPY-C12-based fluorogenic lipase substrate produced strong signal in untreated human milk but low signal after scalding or orlistat treatment, consistent with lipase-dependent activity under our imaging conditions (Figure S3). Together, the exogenous-lipase challenge and endogenous-lipase activity measurements support a model in which it is essential to preserve the MFGM structure to limit lipase access to the TAG core, although assessing the relative contributions of BSSL and LPL to MFGM degradation during storage will require further study.

Despite the physiological importance of lipases in infant fat digestion and gut development, uncontrolled lipolysis during storage leads to accumulation of FFAs associated with rancid flavor. Human milk is enriched in short- and medium-chain fatty acids, such as butyrate (C4), caprylate (C8), and caprate (C10)^31,32^, which contribute to off-odors when released by lipases^5^. Our metabolomic analyses show that the PBF reduces the accumulation of these lipolysis products (Figure 4). In agreement with these findings, sensory evaluation using a blinded forced-choice paradigm demonstrated that PBF-treated samples were more often perceived as similar to fresh milk compared to controls (Figure 5). This finding is significant because changes in stored-milk smell and taste have been linked to infant refusal^4,5^, and the ability of the PBF to preserve sensory acceptability represents a direct, measurable benefit for families relying on stored expressed milk.

In addition to lipolysis, lipid oxidation also contributes to MFG deterioration during storage. PUFAs, although nutritionally important, are especially prone to oxidation^33^. Our oxylipin profiling reveals that PBF-treated samples exhibit reduced accumulation of 9-HODE, 13-HODE, and 12(13)-EpOME (Figure 4). This effect may reflect, in part, the presence of antioxidant components such as vitamins C and E in the formulation. The presence of PUFAs within MFGMs, primarily in the form of glycerophospholipids, suggests the possibility that oxidation and membrane disruption may be linked^34^. Oxidative damage to membrane lipids could weaken the MFGM structure^35^, making the core more accessible to lipases and generating additional PUFAs and diacylglycerols that can further oxidize^36^, reinforcing the importance of preserving the MFGM.

The protective effect of the PBF likely arises from a combination of mechanisms. Pectin is a negatively charged polysaccharide that can chelate Ca^2+^, which has been associated with lipase activity^37,38^. Pectin may also interact with the MFGM to generate an additional protective layer, or it may drive electrostatic repulsion that reduces lipase access to MFGMs. Consistent with preserved MFGM integrity, our proteomics data show that the overall MFGM protein composition is maintained in PBF-treated samples (Figure S4), suggesting that pectin stabilizes the MFGM without displacing resident proteins. This finding may reflect electrostatic interactions between the negatively charged pectin and the positively charged membrane-associated proteins that reinforce the MFGM barrier to lipase access. Although pectin has been reported to exhibit antimicrobial activity in a wide range of organisms^39^, we did not observe significant changes in microbial burden (Figure S6), suggesting that the protective effects observed here are unlikely to be driven by antimicrobial mechanisms.

Several limitations of our studies warrant consideration. As with most human milk research, sample sizes were modest, and larger, more diverse donor cohorts are needed to establish generalizability. Only a single PBF was tested, leaving dose-response relationships unexplored. Because the formulation would be added to milk that infants consume, future studies should also evaluate tolerance, gastrointestinal motility, and microbiome effects following ingestion of pectin-containing human milk. Connections between maternal dietary patterns and formulation efficacy remain to be examined. Despite these limitations, the results reported here establish MFGM destabilization as a measurable outcome of lipid deterioration in stored human milk.

From a practical perspective, our findings suggest that preserving MFGM integrity during storage may improve the longevity and usability of expressed, stored human milk. Milk rejection due to off-flavor is a common challenge for lactating parents and can result in substantial waste of stored milk. A formulation based on pectin, a food-compatible, generally recognized as safe (GRAS) ingredient combined with dietary antioxidants offers a simple approach to limit storage-associated deterioration without altering overall human milk composition. These findings highlight the potential for supplementation of human milk with the PBF as a practical strategy to improve human milk stability during storage.

## Methods

### Human milk collection and donor eligibility

Freshly expressed human milk was collected from 20 lactating donors under a prospective, non-interventional protocol approved by the Sterling Institutional Review Board (IRB ID 12409). Eligible participants were healthy individuals aged 18 to 50 years who were actively breastfeeding a child 1 to 24 months of age, resided in New Jersey, New York, or Pennsylvania, were English-speaking, and were able to provide at least 120 mL of expressed milk without disrupting infant feeding routines using a breast pump. Self-reported exclusion criteria included non-lactating status, participation in another interventional trial, and current diagnosis or treatment for hepatitis B or C, HIV/AIDS, tuberculosis, or herpes simplex virus with active breast lesions. All participants provided electronic informed consent prior to study procedures and sample collection.

Participants provided a single sample of at least 120 mL of freshly expressed milk collected during their usual pumping routine. Milk was transferred into the participant’s usual collection bag or a study-provided sterile bag and labeled with coded participant ID, date, and time of expression. Samples were expressed on-site or delivered by the participant to the laboratory within 2 h of expression. Upon receipt, milk was logged under a coded donor ID and aliquoted into pre-labeled tubes for downstream analyses.

### Formulation preparation

Standardized low-methoxy pectin (∼50% pure pectin based on third-party analysis at Medallion Labs) was prepared as a 4% (w/v) stock solution in deionized water. Due to limited solubility, pectin was dissolved in a 95 °C water bath with continuous agitation for 1 h, then cooled overnight at 4 °C. The following day, the pectin solution was brought back to room temperature. Separately, concentrated solutions of vitamin C (20.2 mg/mL) and vitamin E (1.8 mg/mL) were prepared and mixed at a 1:1 (v/v) ratio. The vitamin mixture was added to the pectin solution at a 1:9 (v/v) ratio. The resulting pectin-vitamin formulation was added to human milk to a final concentration of 0.4% (w/v), corresponding to 0.2% (w/v) pectin.

### Fluorescent staining of the MFGM using Di-4-ANEPPDHQ for confocal microscopy

Di-4-ANEPPDHQ was obtained from Invitrogen (D36802). A 1 mg/mL stock solution was prepared in dimethylsulfoxide (DMSO) and stored at −20 °C. Milk samples were thawed from −80 °C, aliquoted, treated with formulation or an equal volume of water as the control, and stored at either 4 °C or −20 °C for subsequent time-point imaging or the samples were returned to −80 °C for long-term storage. The Di-4-ANEPPDHQ stock solution was diluted 1:1,000 into the milk samples. Stained samples were incubated for at least 30 minutes at room temperature with gentle agitation to allow uniform dye partitioning into the MFGM. For mounting, 50 µL of stained milk was combined with 50 µL of 1% low melting point agarose in PBS, pH 7.4. A 50 µL aliquot of the mixture was placed on a microscope slide, covered with a coverslip, and allowed to solidify for at least 30 minutes before imaging. Fluorescence imaging was performed on a Zeiss LSM880 confocal microscope equipped with a 63× oil-immersion objective. Di-4-ANEPPDHQ was excited using a 488 nm laser at 1% power with a detector gain of 700 and 8× frame averaging. Images of commercial MFGM-supplemented formula (Enfamil NeuroPro) were acquired using the identical laser settings and objective, with Airyscan mode (default processing settings) and 4× frame averaging, to improve signal-to-noise. Images used for radial fluorescence profile analyses in Figure S1 were acquired as z-stacks to avoid bias from selective focal-plane imaging, with 1× frame averaging because residual MFGM movement within the agarose matrix precluded higher averaging without introducing a motion artifact.

Radial fluorescence profiles were generated from individual MFGs by measuring fluorescence intensities from the MFG centers to the maximum fluorescence signals and calculating the center-to-maximum fluorescence ratios. Individual MFG profile measurements were summarized at both the profile and donor-mean levels. Profile-level measurements were modeled with treatment as a fixed effect and donor as a random intercept to quantify image-level distributional shifts. Because individual MFGs are nested within donor samples, treatment consistency was interpreted from donor means; donor-clustered sensitivity analyses were used to evaluate whether profile-level shifts generalized across donors.

### Lipase treatment of milk

Lipase from *Pseudomonas cepacia* (Sigma-Aldrich, 62309; specific activity ∼34 U/mg) and from porcine pancreas (Carolina Biological Supply Company, 872500; specific activity ∼19 U/mg) were used. Following Di-4-ANEPPDHQ staining and incubation of milk samples, lipases were added to a final concentration of 0.02 U/mL for bacterial lipase and 4 U/mL for porcine pancreatic lipase. After gentle mixing, 10 µL of lipase-treated milk was loaded onto microscope slides pre-treated with 1 mg/mL concanavalin A (ConA) by drying the solution between the slide and coverslip to immobilize MFGs. Samples were covered with ConA-coated coverslips separated by double-sided tape spacers to prevent compression. The edges were sealed with clear nail polish to minimize evaporation.

### Endogenous lipase activity imaging

To visualize endogenous lipase activity, untreated human milk, human milk scalded at 82 °C for 10 minutes, and orlistat-treated human milk were incubated for 1 h with a BODIPY-C12-based fluorogenic lipase substrate (EnzChek Lipase Substrate, Invitrogen/Thermo Fisher Scientific, E33955). For orlistat (Ambeed, A52085) treatment, a 10 mM stock solution was prepared in ethanol, then added to human milk to a final concentration of 100 µM, followed by 30 minutes incubation at 37 °C before adding the EnzChek lipase substrate. Samples were imaged by confocal microscopy and substrate-derived fluorescence compared.

### Biochemical profiling of the matched three-donor storage subset

Proteomics, metabolomics, and oxylipin analyses were performed on a matched set of three independent donors analyzed at baseline and after 4 days at 4 °C, and 7, 30, and 90 days at −20 °C. Untreated and PBF-treated samples were analyzed in parallel.

### Proteomic analyses of human milk

Human milk samples (1.75 mL) were subjected to centrifugation at 4 °C for 30 minutes at 4,000 × g to separate the lipid layer from the aqueous phase. The collected aqueous layer was again subjected to centrifugation at 4 °C for 30 minutes at 4,000 × g. The resulting aqueous phase was collected, and 30 µL was subjected to 10-kDa molecular weight cut off (MWCO; Pall Laboratory, Port Washington, New York, USA) filtration to isolate proteins and minimize residual pectin interference. Because pectin co-purifies with proteins, absolute protein recovery was lower in PBF-treated samples than in untreated controls. However, relative protein abundances were highly correlated between matched treatment groups, indicating that the formulation does not broadly alter the human milk proteome despite differences in total yield. Retained proteins were combined with 5 µL of 2 M ammonium bicarbonate (Thermo Scientific, Waltham, MA, USA), reduced with 2 µL of 550 mM dithiothreitol (Promega, Madison, WI, USA) at 50 °C for 50 minutes, alkylated with 4 µL of 450 mM iodoacetamide (Sigma Aldrich, St. Louis, MO, USA) at room temperature for 60 minutes in the dark, and digested overnight at 37 °C with 2.5 µL of trypsin/Lys-C (1 µg/µL, Promega). Trypsin-derived peptides were purified by C18 solid-phase extraction (Biotage, Uppsala, Sweden) and dried by vacuum centrifugation. Dried samples were reconstituted in 100 µL of 3% acetonitrile (Merck, Darmstadt, Germany) with 0.1% formic acid (Sigma Aldrich) and diluted 10-fold. Peptides were analyzed by ultra-performance liquid chromatography on a Waters nanoACQUITY instrument (Waters, Milford, MA, USA) coupled to an Orbitrap Fusion Lumos mass spectrometer (Thermo Scientific). Peptide identification was performed in ProteomeDiscoverer v3.2 against an in-house human milk protein database (374 sequences), and relative protein abundances were compared across storage conditions and treatment groups.

### NMR metabolomics analysis

Targeted aqueous-phase ^1^H-NMR metabolomics was performed on the matched three-donor storage subset, including untreated and PBF-treated samples at baseline and following storage for 4 days at 4 °C or 7, 30, and 90 days at −20 °C. Samples were prepared using established human milk NMR metabolomics workflows^40–42^. Briefly, milk aliquots were thawed on ice and subjected to centrifugation at 16,000 × g for 5 minutes at 4 °C to separate the lipid and aqueous fractions. The aqueous fraction was removed and filtered through Amicon Ultra-0.5 mL 3-kDa MWCO centrifugal filters (Millipore Sigma, Burlington, MA, USA) at 16,000 × g for 50 minutes at 4 °C to remove high-molecular-weight material, including proteins, residual lipids, and pectin. Before sample filtration, filters were washed three times with 500 µL Milli-Q water to remove the glycerol preservative.

For each sample, 207 µL of filtrate was combined with 23 µL of Chenomx internal standard solution containing 5.0 mM 3-(trimethylsilyl)-1-propanesulfonic acid-d6 (DSS-d6), 0.2% NaN_3_, and 99.8% D_2_O. Sample pH was measured and adjusted to 6.8 ± 0.1, and 180 µL of the final mixture was transferred to a 3-mm NMR tube. Samples were maintained at 4 °C until spectral acquisition.

^1^H-NMR spectra were acquired at 25 °C on a Bruker Avance 600-MHz NMR spectrometer equipped with a SampleJet autosampler using a one-dimensional NOESY-presaturation pulse sequence (noesypr1d) for water suppression. Spectra were Fourier transformed, manually phased, and baseline corrected using Chenomx Processor within the Chenomx NMR Suite. Metabolites were assigned and quantified using the Chenomx Profiler by comparison with reference spectral libraries and by normalization to the DSS-d6 internal standard. Metabolite concentrations were corrected for dilution and reported as µmol/L.

A total of 59 polar metabolites were quantified across storage conditions. For the present analysis, glycerol was interpreted as a direct product of triacylglycerol hydrolysis, and butyrate together with the partially overlapping caprate/caprylate resonance region were selected as markers of storage-associated short- and medium-chain fatty acid release. All quantified metabolites are provided in the Source Data.

### Oxylipin analysis by liquid chromatography-tandem mass spectrometry (LC-MS/MS)

LC-MS/MS oxylipin profiling was performed on the above matched set. Analyses included matched free and total measurements where available and focused on free oxylipins, including 13-HODE, 9-HODE, and 12(13)-EpOME^43^.

### Bulk macronutrient and glycerol profiling

Macronutrient milk composition was evaluated by Fourier transform infrared-based compositional profiling on a FOSS MilkoScan FT3 instrument across baseline and selected storage conditions on human milk samples from a broad donor cohort. Analyses focused on fat, protein, and lactose as bulk macronutrient readouts, with glycerol serving as an indicator of storage-associated lipolysis.

### Microbial plate counting

Microbial plate counting was performed on a 10-donor subset from the larger study population. Untreated and PBF-treated human milk samples were plated on blood agar, MacConkey agar^44^, and Sabouraud agar (ASM protocol, https://asm.org/protocols/sabouraud-agar-for-fungal-growth-protocols; accessed 26 Mar 2026) at baseline, after 6 h at room temperature, and after 6 days at 4 °C. Counts are reported as CFU/mL. Sabouraud agar was used as a fungal screen.

### Lipase activity measurement and transcriptomic correlation in an independent cohort

Net lipase activity was measured in n = 200 human milk samples from an independent Mothers and Infants LinKed for Healthy Growth (MILK) Study cohort collected at 1 and 6 months postpartum (1-month n = 53, 6-month n = 147) using a 4-nitrophenyl octanoate (4-NPO; Sigma-Aldrich) lipase activity assay. Total RNA was extracted from the cellular fraction of each sample, and RNA-seq libraries were prepared and sequenced as described previously^45^. Transcript abundances for *CEL* and *LPL* were extracted from the processed expression matrix. The relationship between rank-normalized net lipase activity and rank-normalized *CEL* or *LPL* expression was assessed using a linear regression model including collection time point (1 or 6 months) and the first five principal components of the gene expression matrix as covariates to adjust for known and unobserved factors influencing gene expression. This donor cohort was not the same as those who donated samples used for the MFGM imaging and biochemical profiling described above.

### Olfactory sensory evaluation

To assess whether PBF treatment results in detectable sensory differences, a blinded olfactory evaluation was conducted using commercially available goat milk as a surrogate mammalian system. Sterling Institutional Review Board determined that this evaluation was exempt from IRB review (Sterling IRB ID 15642-JESilpe). Three sample conditions were prepared: a fresh untreated reference, goat milk treated with 4 U/mL lipase (Integrative Therapeutics Lipase Concentrate-HP), and lipase-treated goat milk supplemented with the PBF. Samples were coded and presented in randomized order. Adult panelists (n = 44) first smelled the fresh reference, then smelled the two coded samples and indicated via forced choice which sample smelled most similar to the fresh sample. Panelists rated the perceived similarity of their chosen sample to the fresh reference on a continuous 0–100 scale (0 = not similar, 100 = identical). Responses were collected on an electronic form and analyzed using a two-sided binomial test against the null hypothesis of equal preference (50%) and a Mann–Whitney U test for similarity ratings.

## Acknowledgements

We thank the Mothers and Infants LinKed for Healthy Growth (MILK) Study investigators, staff, and participants for supporting the use of MILK Study samples and associated RNA-seq data. We also thank Sadia Sattar Sultani for contributing to the oxylipin analysis.

## Author contributions

J.E.S., H.K., and B.L.B., conceived the study, designed experiments, analyzed data, and wrote the manuscript. Y.-T.T. contributed Di-4-ANEPPDHQ probe-screening, method development, and microscopy experiments. I.B. conceived the study, designed experiments, and contributed fluorescence microscopy expertise, and data interpretation. A.S. contributed sample collection, experimental execution, and data interpretation. K.E.J. contributed MILK Study RNA-seq and milk-omics data and interpretation. B.J.K. and D.C.D. contributed proteomics analyses and interpretation. C.M.S. contributed NMR metabolomics analyses and interpretation. A.Y.T. contributed oxylipin analyses and interpretation. B.L.B. supervised the work, secured funding, and edited the manuscript. All authors reviewed and approved the manuscript.

## Data availability

Mass spectrometry proteomics data have been deposited with the ProteomeXchange Consortium via the PRIDE partner repository under dataset identifier PXD080793 and DOI https://doi.org/10.6019/PXD080793. Additional data supporting the findings are available in the Supplementary Information and Zenodo at https://doi.org/10.5281/zenodo.21325076.

---

**Figure S1.**
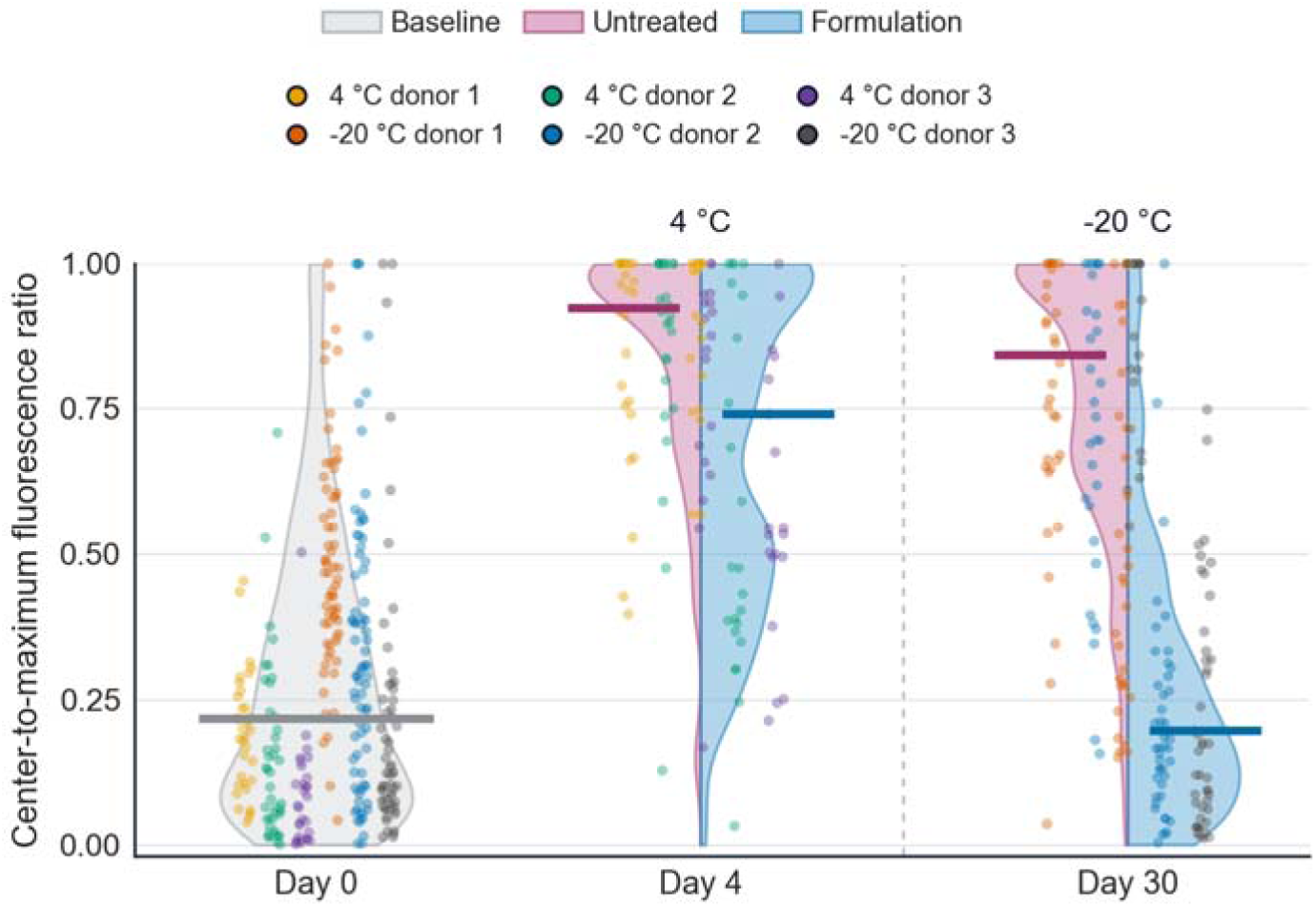
Radial-profile quantitation of Di-4-ANEPPDHQ staining of untreated and PBF-treated milk during storage. Center-to-maximum fluorescence ratios are shown for individual MFGs at baseline (Day 0), after 4 days at 4 °C, and after 30 days at −20 °C. The 4 °C and −20 °C experiments used distinct three-donor sets; Day 0 includes baseline profiles from both experiments. Violin plots show profile distributions; points show individual MFG profiles colored by donor, and horizontal bars indicate medians. PBF-treated aliquots showed lower ratios in two of three donors at 4 °C and in all three donors at −20 °C (donor-level paired tests, P = 0.222 and P = 0.0495, respectively).

**Figure S2.**
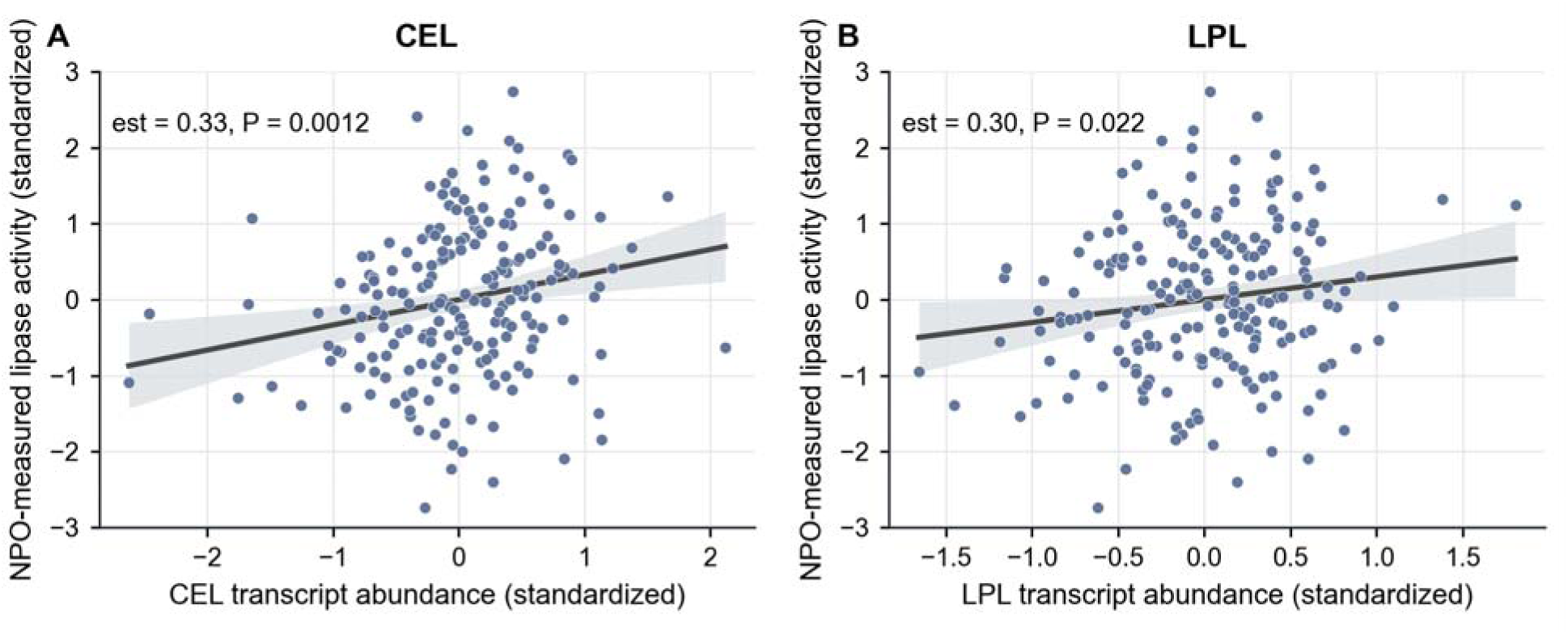
Correlation between lipase transcript abundance and 4-nitrophenyl octanoate (4-NPO)-measured lipase activity in an independent human milk cohort. (A) Standardized *CEL* transcript abundance, encoding BSSL, versus standardized lipase activity (est = 0.33, P = 0.0012). (B) Standardized *LPL* transcript abundance versus standardized lipase activity (est = 0.30, P = 0.022). Each point represents one human milk sample. Samples collected at 1 and 6 months postpartum are shown.

**Figure S3.**
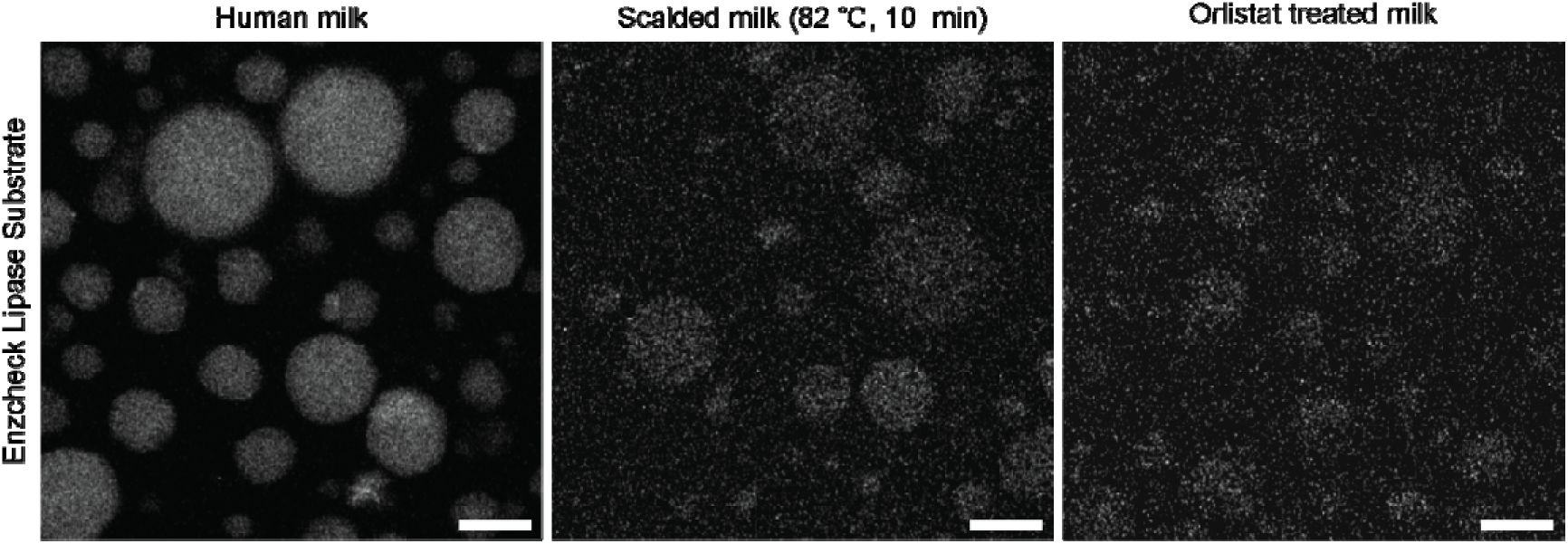
Endogenous lipase activity in human milk. Confocal microscopy images of human milk incubated for 1 h with a BODIPY-C12-based fluorogenic lipase substrate (EnzChek). Left: untreated human milk, middle: scalded milk (82 °C, 10 minutes), right: orlistat-treated milk. Scale bars, 10 µm.

**Figure S4.**
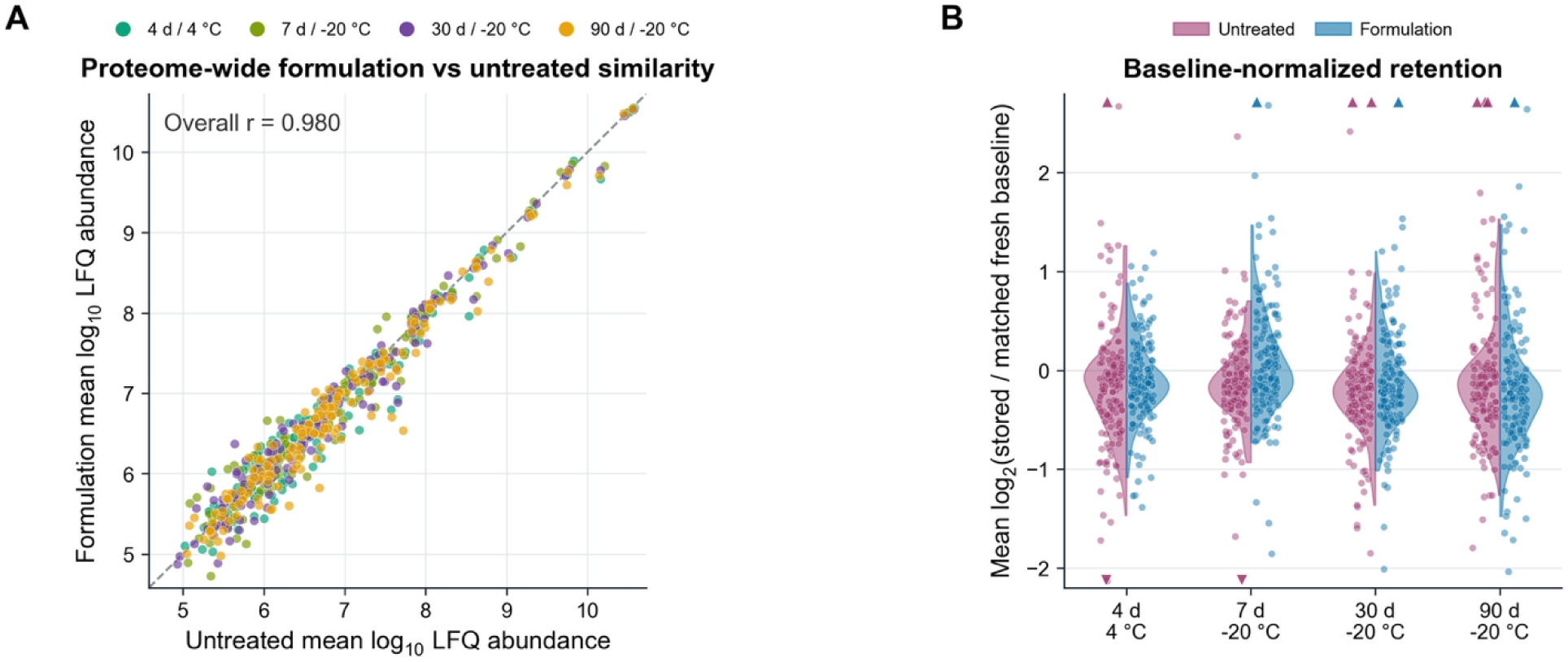
Proteomic comparison of untreated and PBF-treated human milk across storage conditions. (A) Mean log_10_ label-free quantification (LFQ) abundance per protein in PBF-treated versus matched untreated samples, colored by storage condition as designated; the dashed line indicates identity. (B) Baseline-normalized protein retention distributions shown as split half-violins as mean log_2_(stored / matched fresh baseline), with individual protein values overlaid. Colors show untreated and PBF-treated conditions as designated. Boundary triangles indicate values outside the plotted y-axis range; panel B includes all quantified proteins (n = 162).

**Figure S5.**
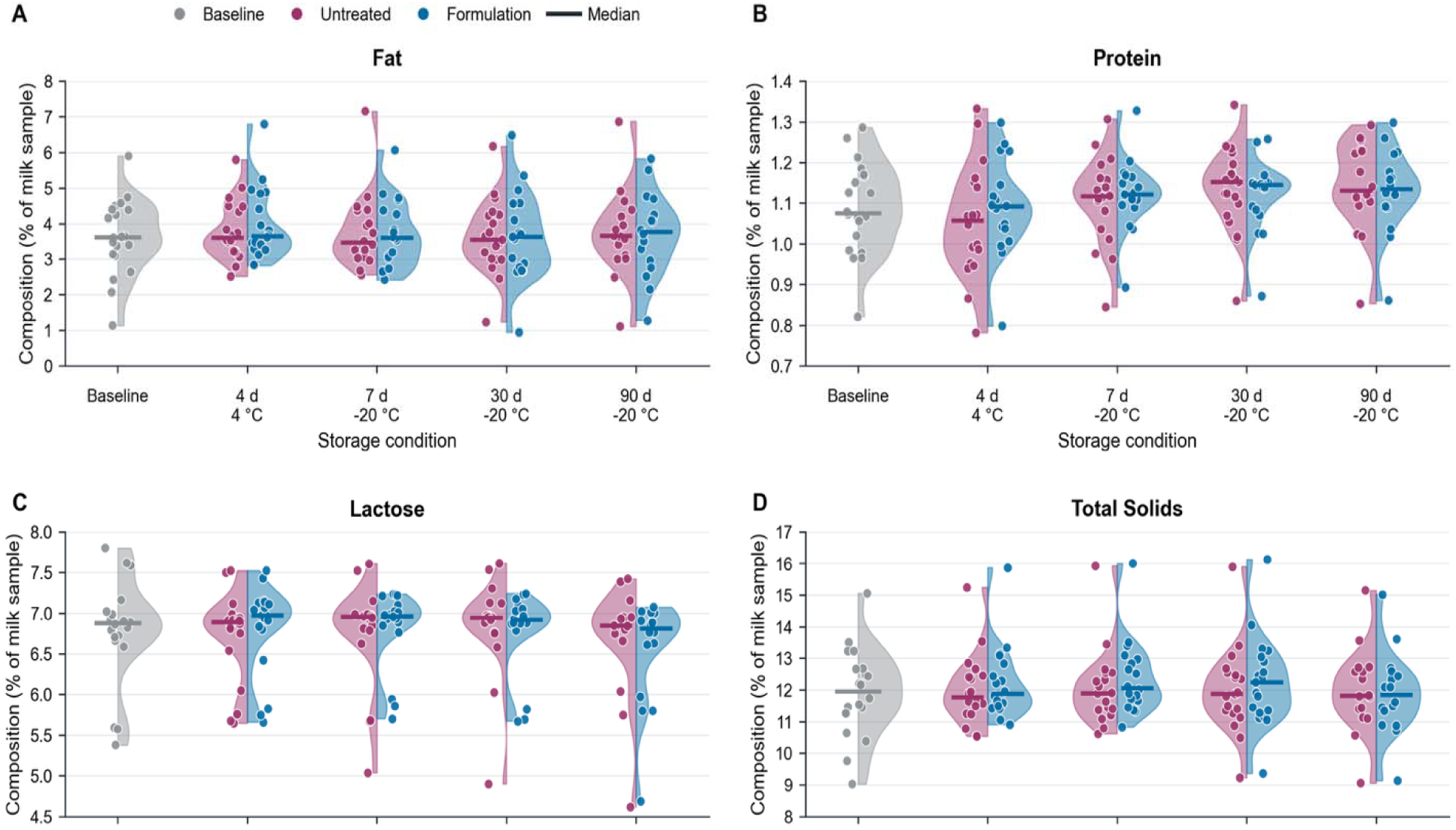
Fourier transform infrared-based bulk macronutrient profiling of untreated and PBF-treated human milk. Panels show Fourier transform infrared (FTIR)-derived fat (A), protein (B), lactose (C), and total solids (D), reported as percent composition of the human milk sample, across the baseline (untreated day-0) and designated storage conditions.

**Figure S6.**
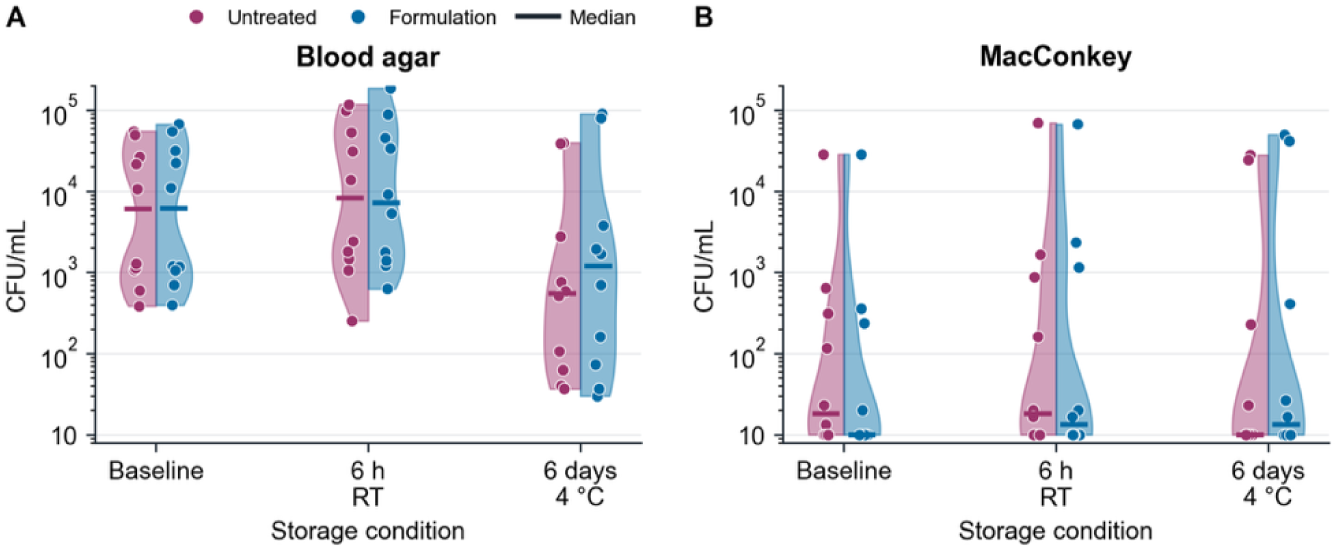
Microbial plate counts for untreated and PBF-treated human milk at 4 °C. Counts on blood (A) and MacConkey (B) agar are shown on log CFU/mL axes. Values below 10 CFU/mL are plotted at 10 CFU/mL for ease of visualization. Sabouraud plates showed no fungal growth (not shown). A 10-donor subset is shown for each growth medium.

